# A universal microfluidic approach for quantitative study of bacterial biofilms

**DOI:** 10.1101/2021.08.24.457583

**Authors:** Yuzhen Zhang, Lingbin Zeng, Yumin Cai, Zhaoyuan Chen, Peng Liu, Luyan Z. Ma, Jintao Liu

## Abstract

Bacteria usually live in densely packed communities called biofilms, where interactions between the bacteria give rise to complex properties. Quantitative analysis is indispensable in understanding those properties. However, current biofilm culturing approaches impose various limitations to these types of analysis. Here, we developed a microfluidic approach for quantitative study of biofilms, which is universal and can be used to culture biofilms of various bacterial species. To demonstrate the advantages of this approach, we present two examples, both of which revealed new biological insights. In the first example, we explored the response of *Escherichia coli* biofilms to exogenous hydrogen peroxide; We found the biofilms gained resistance to H_2_O_2_, but their growth was slowed down due to the metabolic cost of maintaining the resistance; However, under oxygen limitation, H_2_O_2_ can anti-intuitively boost biofilm growth. In the second example, we explored resource retention by *Pseudomonas aeruginosa* biofilms; We observed a fluorescent substance within the biofilm and identified it as the siderophore pyoverdine; We further showed that the extracellular matrix component Psl acted as a retention barrier for pyoverdine, minimizing its loss into the environment and therefore potentially promoting sharing of pyoverdine within the biofilm.

## Introduction

Bacteria usually live in communities, which is beneficial for their survival ^1–3^. Many communities exist on surfaces, where the bacteria secrete extracellular polymeric substances such as proteins, polysaccharides, and extracellular DNA, which enable the bacteria to form dense aggregates ^4^. These aggregates are called biofilms. It was estimated that biofilms accounted for the majority of the bacteria in nature ^5^. In addition to being environmentally important, biofilms are also closely related to our health. For example, they are often the cause of infections. Since biofilms are highly tolerant to antimicrobials and to our immune system, those infections are often persistent and are hard to cure ^6^.

A distinguishing feature of biofilms is spatial heterogeneity ^7^. Due to the dense packing of the bacteria, spatial gradients emerge within biofilms. For example, biofilm periphery is relatively nutrient rich while biofilm interior is nutrient poor; On the other hand, metabolic by-products accumulate at biofilm interior and dissipate toward biofilm periphery. These spatial gradients dictate that bacteria at different locations of the biofilm experience different environments. As a consequence, the bacteria display a broad spectrum of physiological states within a single biofilm, and different regions of the biofilm can have distinct properties ^8–13^. Moreover, recent studies found that interactions between different regions of a biofilm gave rise to emergent collective behaviors that are beneficial to the survival of the bacteria ^14–16^. Therefore, systematic studies on the spatial heterogeneity can help us to gain a deeper understanding of biofilms and their resistance.

We developed a microfluidic approach of culturing biofilms, which facilitates quantitative analysis. In our previous effort, we adapted a commercial microfluidic chip that was designed for yeast single-cell studies to the culturing of *Bacillus subtilis* biofilms ^14^. However, it is unsuitable for many other species of bacteria, especially those of medical importance. The main reason is that many bacteria display strong preference for adhesion, which causes unintended seeding of bacteria at random locations and leads to clogging of the microfluidic chip, hindering long-term tracking of biofilm properties. Here we developed a new microfluidic design from scratch, which overcame the seeding issue and can be applied to all the commonly studied bacterial species. Moreover, the microfluidic chip can be manufactured in a standard microbiology laboratory, and can be easily modified to accommodate the needs of specific experiments. In the following, we will describe our new design, and will also give two examples to demonstrate how our method could be used to reveal new biological insights.

## Results

### Description and characterization of the method

Figure 1a is a schematic diagram of our microfluidic chip. A key component is a specially designed bacteria seeding zone 7 on the side of the biofilm growth chamber 3 (Fig. 1a, marked in red; Methods). During loading, we inject planktonic bacteria culture into the loading port 5; The injection pressure creates a narrow gap at the seeding zone, allowing bacteria to pass through; Some of the bacteria are trapped at the seeding zone, while the rest are flushed out through the waste outlet 4 (Fig. 1b, Methods). In this way, we could plant bacteria specifically at the seeding zone. The trapped bacteria later proliferate into the growth chamber, where fresh medium is continuously supplied. By secreting extracellular matrix, the bacteria are able to form a stable and densely packed biofilm in the presence of flow (Fig. 1c). Figure 1d shows time-lapse images of an *E. coli* biofilm, which emerged out of the seeding zone ^~^8 hours after loading, and later grew into a round shaped biofilm containing millions of bacteria (Supplementary Video 1).

**Fig. 1:**
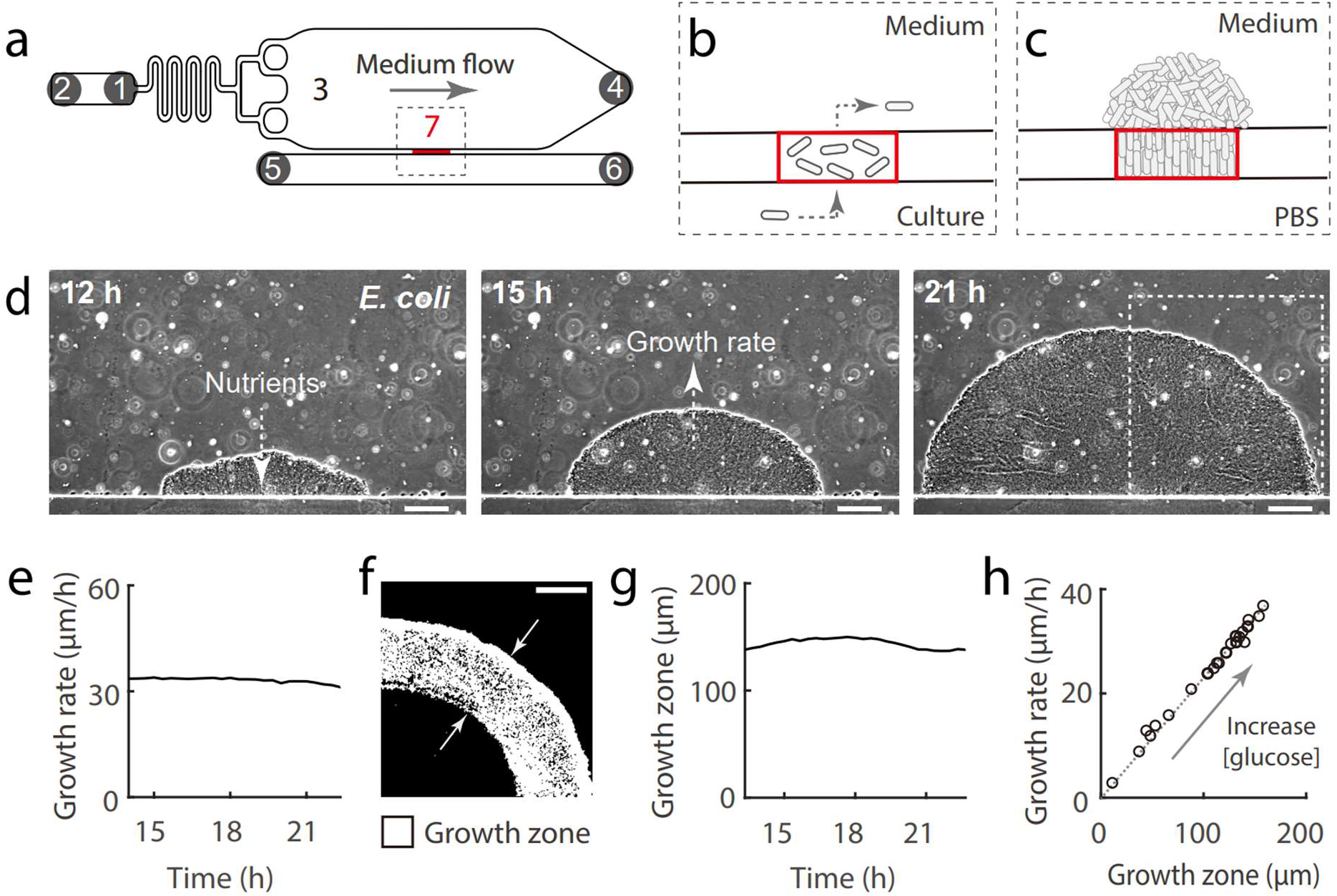
Microfluidic design and growth of *E. coli* biofilm. **a**, Schematic diagram of the microfluidic device: 1) medium inlet; 2) medium exchange port; 3) growth chamber; 4) waste outlet; 5 & 6) bacteria loading channel and ports; 7) bacteria seeding zone (marked in red). **b**, Schematic diagram of the bacteria loading process: The dashed rectangular region in **a** is shown; Planktonic culture of bacteria were injected into the loading channel; Some of the bacteria were squeezed into the seeding zone (marked in red) by the injection pressure, and those that were squeezed through were flushed out through the waste outlet. **c**, Schematic diagram of a biofilm growing out of the seeding zone. **d**, Phase contrast images of a growing *E. coli* biofilm; Three representative time points are shown; Scale bar, 100 μm. **e**, The growth rate of the biofilm shown in **d**. **f**, Differencing analysis of the phase contrast images (Methods) revealed zone of growth in biofilm; The dashed rectangular region in **d** is shown; Scale bar, 100 μm. **g**, The width of the growth zone shown in **f**. **h**, Biofilm growth rate was proportional to the width of the growth zone, and the two were determined by glucose concentration.

We used a thin growth chamber (6 μm thick). In comparison, the radius of the biofilm is on the order of 10^2^-10^3^ μm. Therefore, the biofilm is a pancake-like colony with uniform thickness (Fig. 1d). A major advantage of this design is that the spatial and dynamical properties of the biofilm could be easily quantified. We analyzed the growth of the *E. coli* biofilm, and found that its radius increased at a constant rate (defined as biofilm growth rate) (Fig. 1e). Image analysis revealed that growth was mainly from the periphery region of the biofilm (Fig. 1f, Methods), and the depth of the growth zone was constant over time (Fig. 1g). The biofilm was cultured with M63B1, a defined minimal medium using glucose as the carbon source ^17^. Since nutrients penetrate the biofilm through diffusion (Fig. 1d), and are consumed by the densely packed bacteria during the process, there is an inward gradient of nutrients in the biofilm. Therefore, the growth arrest at biofilm interior should be due to nutrient limitation. We found the depth of the growth zone increased with glucose concentration (Supplementary Fig. 1a), suggesting glucose was the limiting nutrient. Interestingly, we also observed a proportional relationship between biofilm growth rate and the depth of the growth zone (Fig. 1h). Finally, we confirmed that the biofilm growth rate was independent of the chamber thickness (Supplementary Fig. 1b), which gives flexibility to the design of the microfluidic chip.

Another important aspect is the maintenance of oxygen gradient in the biofilm. There are steep O_2_ gradients in natural biofilms, and biofilm interior is usually hypoxic. Oxygen gradient plays important roles in biofilm physiology ^11,18^. Therefore, we need to preserve it in our microfluidic chip. However, a technical issue is that the polydimethylsiloxane (PDMS) material commonly used in microfluidics is gas permeable, which leads to vertical diffusion of O_2_ from the ambient environment into the growth chamber (Supplementary Fig. 2a), causing loss of O_2_ gradient. To overcome this problem, we sealed the PDMS with cover glass and epoxy resin (Fig. 2a), both of which are gas impermeable; The glass covers the chamber portion of the PDMS, so that the biofilm could be observed with microscope; The epoxy resin covers the rest of the device, which also provides mechanical support to the tubes. In addition, we made the PDMS layer as thin as possible (1 mm, further decreasing the thickness would cause collapse of the growth chamber), which minimizes lateral diffusion of O_2_ from the flowing medium through PDMS into biofilm interior. To test the effectiveness of this solution, we imbedded the oxygen probe PtTFPP into the PDMS. The fluorescence of the probe is quenched by O_2_, which can be used as a measure of O_2_ availability ^19^. Results confirmed that an O_2_ gradient was generated by the biofilm (Fig. 2b). To test whether the biofilm interior was hypoxic, we utilized the fluorescent protein YFP, which requires O_2_ to maturate into its fluorescent form ^20^; We constructed a YFP expressing E. *coli* strain driven by the *ptsG* promoter; The promoter is activated by glucose limitation and therefore the YFP would mainly be expressed at biofilm interior ^21^. Biofilms formed by this strain only showed background fluorescence in our microfluidic chip (Fig. 2c and d, at 34 h). We reasoned that the YFP protein was expressed at biofilm interior but did not maturate due to lack of oxygen. Indeed, biofilms cultured without oxygen limitation displayed the expected fluorescence distribution (Supplementary Fig. 2b and c, at 34 h). To confirm the existence of immature YFP protein, we inhibited the synthesis of new YFP using the antibiotic tigecycline (Fig. 2d, Supplementary Fig. 2c) and then fed air through the loading channel (Fig. 1a, Methods); YFP fluorescence immediately emerged upon feeding air (Fig. 2c and d). Finally, we confirmed that YFP could mature at the periphery region of the biofilm without feeding air (Supplementary Fig. 3). These results showed that the biofilm periphery had access to oxygen, while the biofilm interior was hypoxic.

**Fig. 2:**
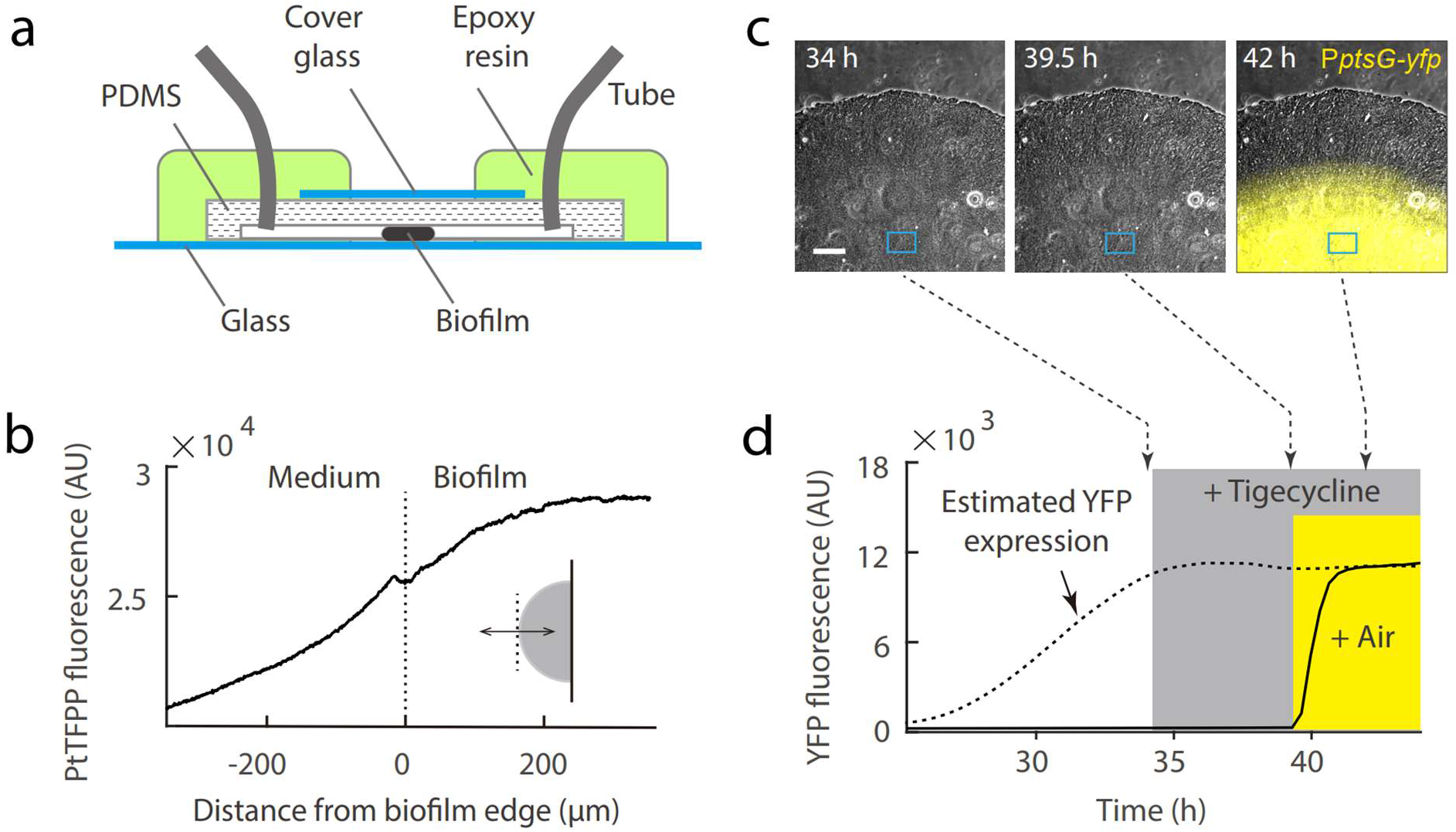
Maintenance of oxygen gradient. **a**, Side view of the microfluidic chip. **b**, Spatial profile of PtTFPP fluorescence over an *E. coli* biofilm; The double arrow in the inset illustrates the line along which the profile was measured. **c**, Time-lapse images of an *E. coli* biofilm; Composite of phase contrast and YFP (shown in yellow) channels; The YFP channel reveals the activity of the *ptsG* promoter; Scale bar, 100 μm. **d**, YFP fluorescence intensity at biofilm interior (the region marked by the rectangle in **c**; The solid line shows the measured fluorescence value; The dashed line shows the estimated level of YFP protein based on the dashed line in Supplementary Fig. 2c; The shaded regions represent the durations of tigecycline (2 μg/ml, shown in gray) and air (shown in yellow) treatments respectively; The dashed arrows indicate the time when the snapshots in **c** were taken.

### Biofilm resistance to oxidative stress

To demonstrate how our method could be used to study biofilms, here we present two examples. The first example is on biofilm resistance. Biofilms are known to show high phenotypic resistance to environmental stresses ^6,22,23^. We investigated the response of E. *coli* biofilms to exogenous hydrogen peroxide, which induces oxidative stress in bacteria and causes DNA damage ^24^. Biofilm growth rate quickly dropped to zero upon switching to medium containing 0.03% H_2_O_2_ (Fig. 3a). However, the biofilm resumed growth after less than 6 hours, showing acquisition of resistance. We monitored the oxidative stress within the biofilm using the reactive oxygen species (ROS) probe DCFH-DA; It showed that H_2_O_2_ was able to affect the periphery region of the biofilm, but had no significant effect on the interior region (Fig. 3b). In *E. coli*, the primary defense machinery against H_2_O_2_ is catalase, which is an enzyme that degrades H_2_O_2_ into H_2_O and O_2_ ^24^. Therefore, we monitored the promoter activity of the catalase gene *katG* using the fluorescent protein mRFP. We found that *katG* was mainly expressed in a narrow region next to the stressed biofilm periphery (Fig. 3b). The narrow size of the catalase band suggested that it was an effective barrier in blocking H_2_O_2_ penetration. Since there was no significant effect on the interior region of the biofilm, we wondered why it did not recover to its pre-H_2_O_2_ growth rate. We speculated that the presence of H_2_O_2_ in the medium imposed a metabolic burden on the biofilm; Specially, although catalase does not consume energy when degrading H_2_O_2_, its lifetime could be limited; Therefore, in order to maintain the H_2_O_2_ barrier, the biofilm had to constantly produce new catalase. Consistent with our speculation, when we removed glucose from the medium, the catalase barrier quickly disappeared and the DCFH-DA band spread toward the biofilm interior (Fig. 3c, Supplementary Fig. 4).

**Fig. 3:**
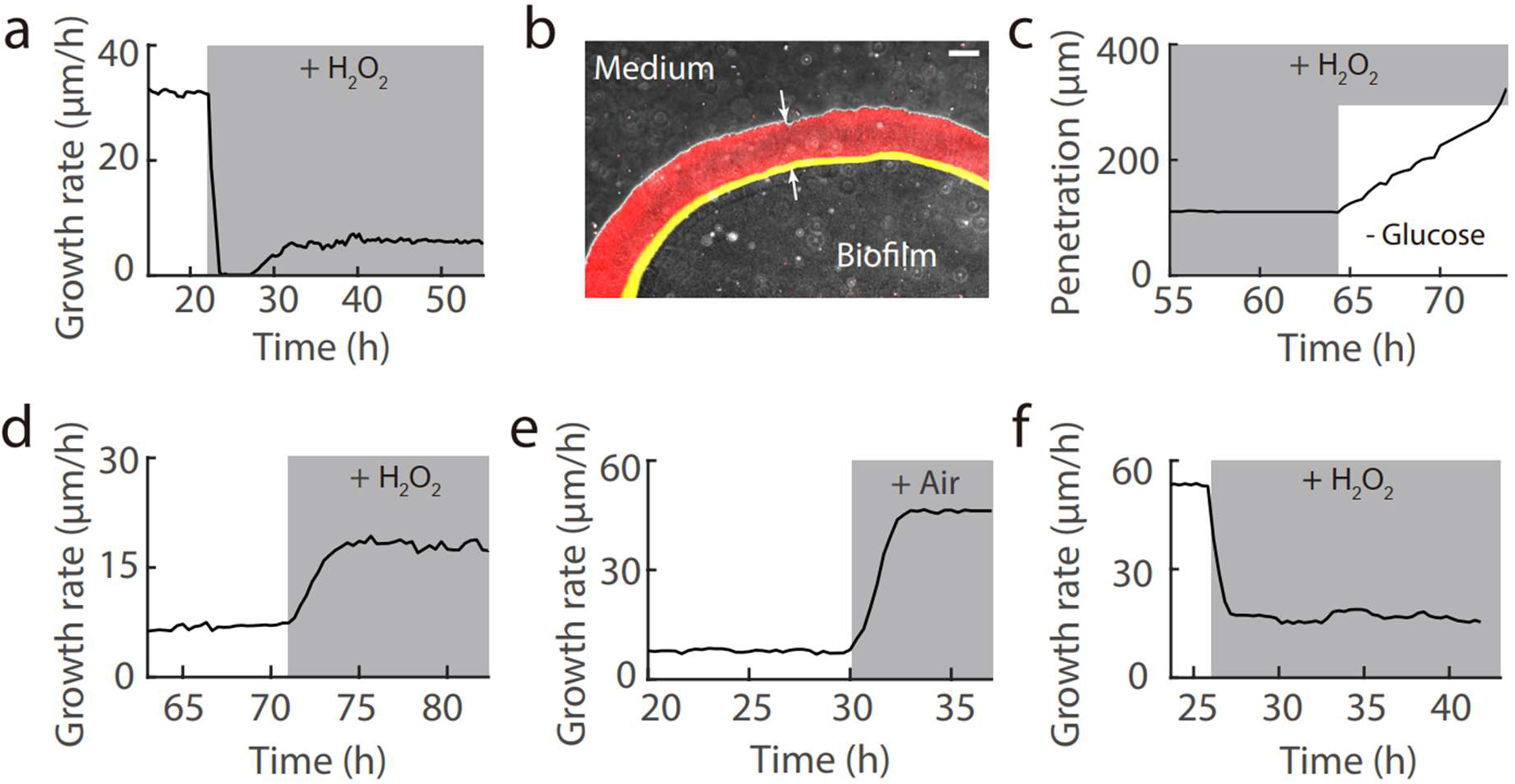
Resistance of *E. coli* biofilm to exogeneous H_2_O_2_. **a**, Biofilm growth rate before and after switching to medium containing 0.03% H_2_O_2_. **b**, Snapshot of biofilm after forming resistance to H_2_O_2_; Composite of phase contrast, YFP (DCFH-DA fluorescence, shown in red), and RFP (*PkatG-mrfp* fluorescence, shown in yellow) channels; The arrows indicate the region of biofilm penetrated by H_2_O_2_; Scale bar, 100 μm. **c**, Depth of H_2_O_2_ penetration before and after removing glucose from the medium. **d-f**, Biofilms cultured with modified M63B1 medium – using glycerol (44 mM) instead of glucose as the carbon source. **d**, Biofilm response to 0.03% H_2_O_2_. **e**, Biofilm response to supplementation of air through the loading channel. **f**, Biofilm response to 0.03% H_2_O_2_, cultured without oxygen limitation (Supplementary Fig. 2a).

We wondered whether external stress always slowed down biofilm growth. Surprisingly, when we used glycerol instead of glucose as the carbon source, H_2_O_2_ counterintuitively boosted biofilm growth (Fig. 3d). The increase of growth rate suggested that H_2_O_2_ might relieved a bottleneck for growth. We wondered whether the bottleneck was oxygen, as degradation of H_2_O_2_ by the catalase barrier could provide additional oxygen to the biofilm. Consistent with our speculation, when we fed air through the loading channel, biofilm growth rate was significantly increased (Fig. 3e); In contrast, doubling the concentration of the carbon source glycerol or the nitrogen source NH_4_^+^ did not have noticeable effect (Supplementary Fig. 5); Finally, when we cultured the biofilm without oxygen limitation (Supplementary Fig. 2a), the growth was suppressed by H_2_O_2_ (Fig. 3f). It is conceivable that this anti-intuitive boosting of growth by stress is not limited to H_2_O_2_. For example, in addition to concentration, diffusion is another important factor affecting nutrient availability inside biofilm; Therefore, one is likely to boost growth if the stress increased the diffusion or transport of the limiting nutrient into the biofilm.

### Resource retention by biofilm

Our method is universal and can be applied to bacteria of various species. In addition to E. *coli*, we also cultured biofilms with *Salmonella typhimurium, P. aeruginosa, Klebsiella pneumoniae, B. subtilis, Staphylococcus aureus, Enterococcus faecium*, and *Mycobacterium smegmatis* (Supplementary Fig. 6), which cover Gram-negative, Gram-positive and mycobacteria. As the second example, here we show our findings on *P. aeruginosa* biofilms. We observed autofluorescence at the interior region of the biofilm under the CFP channel (Fig. 4a). To identify the source of this fluorescence, we took advantage of a well-known behavior of *P. aeruginosa* biofilms – dispersion ^25^. In our experiments, when the biofilm reached the size of ^~^240 μm, the interior cells escaped through an opening at biofilm periphery, leaving behind a hollow shell (Fig. 4a, Supplementary Video 2); Accompanying the dispersion was the simultaneous disappearance of the fluorescence, suggesting loss of the fluorescent substance into the environment; The biofilm later resealed the opening at its periphery, and the fluorescence re-emerged; Interestingly, we observed fluorescence in the hollow region of the resealed biofilm (Fig. 4a, at 13h 10m). These results suggested that the fluorescent substance was extracellular and was diffusible.

**Fig. 4:**
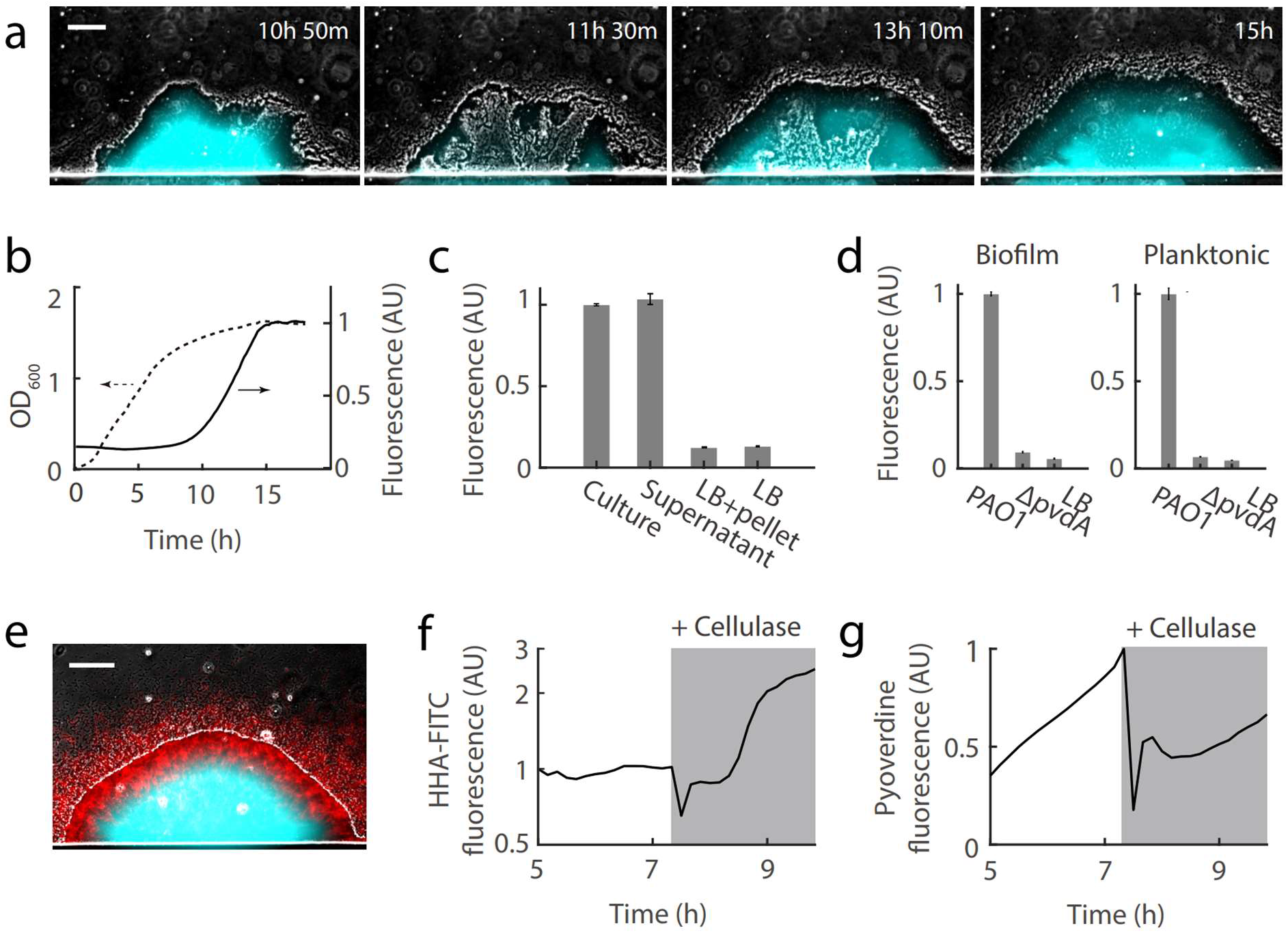
Resource retention by *P. aeruginosa* biofilm. **a**, Time-lapse images of a *P. aeruginosa* biofilm; Composite of phase contrast and CFP (shown in cyan) channels; Scale bar, 100 μm. **b**, Optical density and fluorescence intensity (CFP channel) of planktonic bacteria culture. **c**, Fluorescence intensities of stationary phase planktonic bacteria culture, its supernatant after centrifugation, its resuspended pellet after centrifugation, and the resuspension medium LB; Error bars represent standard deviation, n = 3 biological replicates. **d**, Fluorescence intensities of biofilm (interior region) and stationary phase planktonic cultures with or without *pvdA* knockout. **e**, Staining of biofilm by the fluorescent dye HHA-FITC (shown in red, 100 ug/ml), which binds specifically to the extracellular matrix component Psl; Scale bar, 100 μm. **f**, HHA-FITC fluorescence at biofilm periphery before and after switching to medium containing cellulase (84 U/ml, Sigma C2730-50ML). **g.** Pyoverdine fluorescence at biofilm interior.

Since the fluorescence was most intense at biofilm interior, we suspected it might be related with resource limitation. Using planktonic culture, we confirmed that the autofluorescence emerged when the bacteria entered stationary phase (Fig. 4b). Centrifugation of the stationary phase culture showed that the fluorescence was from the supernatant (Fig. 4c), which confirmed the abovementioned observation that the fluorescent substance in the biofilm was extracellular. Among the substances that *P. aeruginosa* secretes, pyoverdine was known to be fluorescent ^26^. Spectroscopic analysis showed that the excitation and emission spectrum of the stationary phase supernatant was consistent with that of the pyoverdine (Supplementary Fig. 7). Finally, when we knocked out the pyoverdine synthesis gene *pvdA*, the autofluorescence in the biofilm and in the planktonic culture was reduced to the background level (Fig. 4d), confirming the identity of the fluorescent substance as pyoverdine.

Pyoverdine is a siderophore, which is secreted by the bacteria to chelate irons from the environment. Since the process requires re-uptake, uncontrolled loss of pyoverdine into the environment would be energetically inefficient. Therefore, we wondered whether the biofilm was able to confine pyoverdine within its boundary and limit the loss to the flowing medium. A potential mechanism for retention is through the extracellular matrix ^4,27,28^. Ideally, the matrix should not limit diffusion/sharing of pyoverdine within the biofilm. One matrix component that satisfies this criteria is the polysaccharide Psl, which is mainly distributed at biofilm periphery ^12,29^. Therefore, we wondered whether Psl limited the loss of pyoverdine. First, we stained the biofilm using the fluorescently labeled lectin HHA-FITC, which binds specifically to Psl ^29^. Consistent with previous results, fluorescence imaging showed that Psl was mainly distributed at the periphery region of the biofilm (Fig. 4e, Supplementary Fig. 8a); Interestingly, it extended beyond the boundary of the biofilm, which might be trails deposited by the planktonic cells released from the biofilm ^30^. We then treated the biofilm with cellulase – an enzyme that can hydrolyze Psl ^29^, both the HHA-FITC and the pyoverdine fluorescence rapidly dropped to a lower level (Fig. 4f and g, Supplementary Video 3); Cellulase did not affect pyoverdine fluorescence in planktonic culture (Supplementary Fig. 8b), ruling out quenching of pyoverdine. In contrast, treatment of the biofilm with DNase degraded the extracellular DNA – another important component of the extracellular matrix, but did not have effect on pyoverdine fluorescence (Supplementary Fig. 9). These results suggested that Psl limited the loss of pyoverdine into the flowing medium. Finally, we notice that the cellulase treatment eventually boosted Psl level in the biofilm, which is consistent with previous findings that the bacteria could sense the loss of Psl and increase its synthesis via the master regulator c-di-GMP ^26^.

## Discussion

A commonly accepted method of studying biofilms is the flow cell approach, which utilizes a macro fluidic chamber and the biofilms are attached to the chamber wall ^31^. It mimics biofilms in flowing environments, which represents a major fraction of natural conditions ^5^. Due to the large size of the chamber, biofilms formed in flow cell are usually three-dimensional. In contrast, in our microfluidic approach, we made the growth chamber only a few micrometers thick, so that the biofilms are semi-two-dimensional. Despite the difference, our approach is compatible with the flow cell approach. In fact, the biofilm in our microfluidic device is equivalent to a slice of the biofilm in the flow cell. This is because essential properties such as extracellular environments and cellular states are determined by the distance from biofilm surface instead of lateral position. Indeed, we were able to reproduce phenomena observed in flow cells, such as the dispersion of *P. aeruginosa* biofilms and the spatial distribution of Psl (Fig. 4a and e) ^29^. Furthermore, we were able to perform quantitative measurements on spatiotemporal dynamics. Quantitative analysis had been uncommon in the biofilm research community, and a major obstacle was the morphological complexity of the biofilms in the commonly used culturing approaches ^32,33^. By simplifying the biofilm, we were able to eliminate this obstacle. It is true that morphological complexity could play a role in natural settings. Here we have captured a key feature of natural biofilms – spatial heterogeneity. Therefore, findings based on our approach should be generalizable to more complex scenarios.

Quantitative studies of bacterial communities have mainly been limited to small population size ^21,34–37^. However, some important properties are emergent ones that require a minimum population size ^14–16,38,39^. It has been challenging to culture large bacteria communities in microfluidics, as the fluidic chambers and channels are prone to clogging by the bacteria ^40^. Here we are able to culture bacterial communities containing over a million cells for extended period of time (2-5 days), and maintain precise control over the growth environment throughout the entire duration. The key to our success is a specially designed mechanism for the controlled seeding of bacteria at the micro scale. By separating the loading channel from the growth chamber and only seeding bacteria to the designated cell trap, we avoided the main cause of clogging – random adhesion of bacteria in the growth chamber. The controlled seeding also leads to higher reproducibility of biofilms between different experiments, which is essential for quantitative analysis. Since the seeding mechanism does not rely on the properties of the bacteria, our approach is universal and can be used to culture all the commonly studied bacterial species and potentially many more.

Our microfluidic approach is easy to implement. The entire process – from design to soft lithography to chip making – could be carried out in a standard microbiology laboratory (Methods). Yet, it is also very flexible, and could be extended to study more complex bacterial communities. For example, one could use it to study interactions between biofilms of different species (Supplementary Fig. 10). Finally, in addition to combining it with microscopic observation, we have realized its potential to achieve more, and we are currently taking advantage of modern molecular biology tools to conduct systems levels investigations on bacterial communities.

## Supporting information

Time-lapse video of an E. coli biofilm

Time-lapse video of a P. aeruginosa biofilm

Time-lapse video of a P. aeruginosa biofilm

## Acknowledgements

We thank Jing-ren Zhang and Babak Javid for kindly providing bacterial strains and technical supports, thank Jing-ren Zhang for helpful comments on the manuscript, thank members of the Liu Lab for numerous discussions. JL was supported by the Tsinghua University Independent Research Program (20197030008) and the Tsinghua-Peking Center for Life Sciences, YZ was supported by the National Nature Science Foundation of China (21908129), Chinese Postdoctoral Science Foundation (2018M631481), and the Tsinghua-Peking Center for Life Sciences.

## Author Contributions

YZ and JL designed the study, YZ and JL designed the microfluidic chip, YZ, LZ, YC and ZC performed the experiments, YZ, LZ, YC and JL analyzed the data, YZ and JL wrote the manuscript, all authors discussed the manuscript.

## Data availability

The data that support the findings of this study are available from the corresponding author upon reasonable request.

## Methods

### Microfluidic chip fabrication

The microfluidic chip was designed and manufactured in-house. A scaled drawing of the microfluidic design is shown in Supplementary Fig. 11. The master mold of the chip was fabricated using the Maskless Mold Fabrication System from BlackHole Lab (France), and performing lithography on glass slides using photoresist from Microchemicals (AZ4562). The microfluidic chips were made with polydimethylsiloxane (PDMS) and glass slides. The PDMS part was made by pouring 10:1 (v/v) mixture of Sylgard 184 elastomer and curing agent (Dow Corning, USA) on the master mold, and curing the mixture for 2 hours in an oven at 80 °C. The cured PDMS was carefully peeled off from the mold and punched with holes at the inlet and outlet ports. Then we pasted a narrow (0.5-1 mm) strip of 3M Scotch Magic tape to a designated location of the PDMS (on the barrier between the growth chamber and the loading channel, and perpendicular to the barrier), and treated the PDMS with plasma for 2 mins (SoftLithoBox, BlackHole Lab, France). After the plasma treatment, the tape was quickly removed, and the PDMS was immediately bounded with a glass slide or cover glass (depending on the magnification of microscopic observations later on). The tape prevented the plasma from reaching the covered spot, therefore, creating an unbounded region between the growth chamber and the loading channel, which could be used for bacteria seeding. Finally, we bounded the top surface of the PDMS (middle portion) with cover glass, plugged the inlet and outlet ports with syringe needles (20G, 0.91 mm OD × 0.61 mm ID) and PTFE Tubing (1/16″ OD × 1/32 ID), and sealed the rest of the PDMS with epoxy resin (Fig. 2a).

To visualize the spatial gradient of oxygen (Fig. 2b), we embedded the oxygen sensor PtTFPP (Frontier Scientific Inc., USA) in PDMS: The PtTFPP was dissolved in toluene and thoroughly mixed with PDMS before it was cured; Then the mixture was poured on the mold, and toluene was allowed to evaporate while the PDMS was cured. The final PtTFPP concentration was 1 mg/ml.

### Bacterial Strains

The bacterial strains used in this study are *Escherichia coli* BW25113, *Salmonella typhimurium* ATCC 14028, *Bacillus subtilis* NCIB 3610, *Pseudomonas aeruginosa* PAO1 (Δ*paaP*), *Klebsiella pneumoniae* ATCC BAA-1144, *Staphylococcus aureus* RN4220, *Enterococcus faecium* ATCC 35667, and *Mycobacterium smegmatis mc^2^*-155 (Supplementary Table 1).

#### E. coli strains

Promoters of *E. coli* strains (*PptsG* and *PkatG*) were amplified from wild-type *E. coli* BW25113 genomic DNA; The *PtetR* sequence was synthesis by Genewiz (China). The promoter sequences were amplified by PCR and were then introduced to a plasmid containing fluorescent protein coding sequence to generate promoter activity reporters. Plasmids were constructed using the Gibson assembly method and transformed into DH5α competent cells (weidi, China). DNA purification and isolation of plasmids were performed using reagents from Omega Bio-Tek (USA). PCR reactions were carried out using Q5 High-Fidelity DNA polymerase (NEB, UK).

#### *P. aeruginosa* strains

Most of the experiments were based on the PAO1 Δ*paaP* stain, which was less prone to dispersion than wild-type PAO1 and more suitable for the current study; We have verified that *paaP* knockout does not affect the production of the fluorescent substance (data not shown). The *pvdA* deletion strain were constructed by an unmarked, non-polar deletion strategy as previously described (*Environ. Microbiol. Rep*. **10**, 583–593, 2018).

### Bacteria loading and biofilm culturing

On the day before the experiment, bacteria from −80 °C glycerol stock were streaked on LB agar plates and incubated at 37 °C overnight. On the next day, a single colony was picked from the plate and inoculated to 5 ml of LB broth in a 50 ml conical tube, and incubated at 37 °C in a shaker. After 12 h of incubation, the culture was centrifuged at a relative centrifugal force of 7000 g for 3 min, and then the pellet was re-suspended in 0.5 ml biofilm medium and used for loading into microfluidics. The biofilm medium for *E. coli, B. subtilis, S. typhimurium, K. Pneumoniae, E. faecium* and *M. smegmatis* was M63B1 (100 mM KH_2_PO_4_, 15 mM (NH_4_)_2_SO_4_, 0.8 mM MgSO_4_, 3 μM vitamin B1, 22 mM glucose, adjusted to pH 7.4 with KOH) ^17^; The medium for *P. aeruginosa* was LB without sodium chloride (LBNS) ^29^; The medium for S. *aureus* was TSB.

The microfluidic chip was controlled using the Elveflow OB1 Mk3 Pressure Controller (ELVESYS, France), which drives medium flow with pressurized air. At first, the medium was pumped into the growth chamber from ports 1 and 5 (Fig. 1a) simultaneously, with ports 2 and 6 closed. The pressure was maintained at 15 psi until all the air bubbles disappeared, which usually took 5-10 mins. Then port 5 was switched to planktonic bacteria culture with a pressure of 10-20 psi, while port 1 was maintained at 15 psi; Once bacteria started to enter the growth chamber, the pressures to ports 1 and 5 were adjusted to 1-5 psi and 1-2 psi respectively. After ^~^1000 bacteria cells were trapped at the seeding zone, we turned off the pressure to port 5, and adjusted the pressure to port 1 to 3 psi. Finally, we opened port 6 and washed the loading channel by injecting PBS to port 5 for 30 mins (3 psi). During the rest of the experiment, ports 2, 5 and 6 were kept closed, and port 1 was fed with medium under a constant pressure of 3 psi, and the temperature was kept at 37 °C. To perform perturbations (e.g., adding H_2_O_2_ or antibiotic), port 1 was switched to the corresponding new medium, and port 2 was briefly opened until the residues from the previous medium was completely flushed out and the new medium reached port 1 (this operation allowed fast switching of medium). Normally, the biofilm acquires oxygen from the flowing medium; To supply additional oxygen to the interior region of the biofilm (Fig. 2c-d and Fig. 3e), air was pumped into port 5, with port 6 kept open, and the air could then diffuse into the biofilm from the loading channel; To stop the additional oxygen supply, the loading channel was refilled with PBS and ports 5 and 6 were closed.

### Time-lapse microscopy

The biofilms were observed with phase contrast and fluorescence microscopy. The microscope used was Olympus IX83 (Japan) with Andor’s Zyla 4.2 sCMOS camera (UK). To image the entire biofilm, 10X objective lens was used in most of the experiments. Images were taken every 3 s – 20 min, depending on the question.

### Data analysis

ImageJ (National Institutes of Health, USA) and MATLAB (MathWorks, USA) were used for data analysis. To detect region of expansion in a biofilm, we performed image differencing on snapshots of the biofilm from time-lapse microscopic images; Specifically, we compared the difference between two consecutive phase contrast images by calculating the absolute difference between corresponding pixels in the two images; Since expansion leads to pixel level changes and no expansion leads to little change, image differencing could reveal region of expansion in the biofilm.

### Planktonic culture experiments

The culturing and measurement of planktonic *P. aeruginosa* (Fig. 4b-e) were performed using the SPARK microplate reader (TECAN, Swiss). The bacteria were grown overnight in the LBNS medium in a 37°C shaker. On the next day, 4 μl of the culture were inoculated to 200 μl of fresh LBNS and incubated in 96-well plates. Optical density (600 nm) and fluorescence intensity (excitation 430 nm, emission 480 nm) were recorded every 10 min. The excitation and emission spectrum of the supernatant (Fig. 4d) were also measured using the plate reader.

---

**Supplementary Fig. 1:**
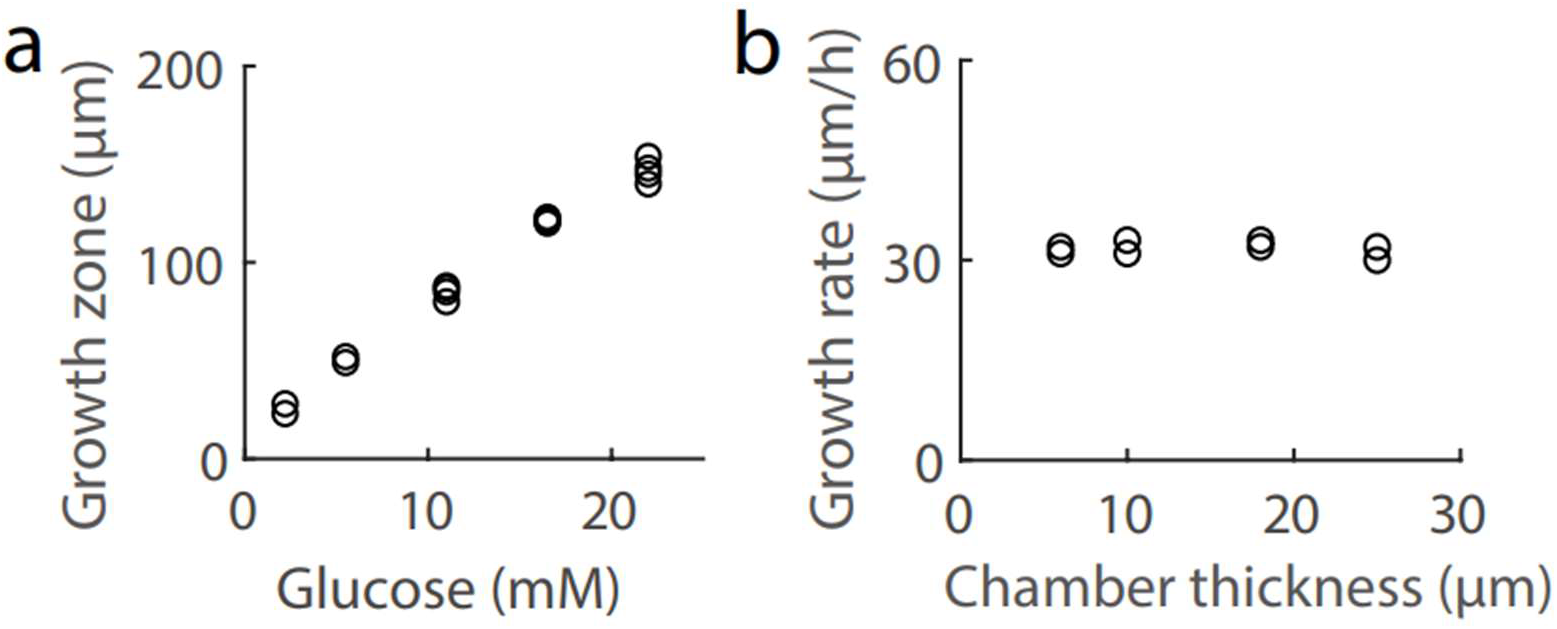
Growth of *E. coli* biofilm. **a**, The depth of the growth zone increased with glucose concentration. **b**, Biofilm growth rate was independent of chamber thickness.

**Supplementary Fig. 2:**
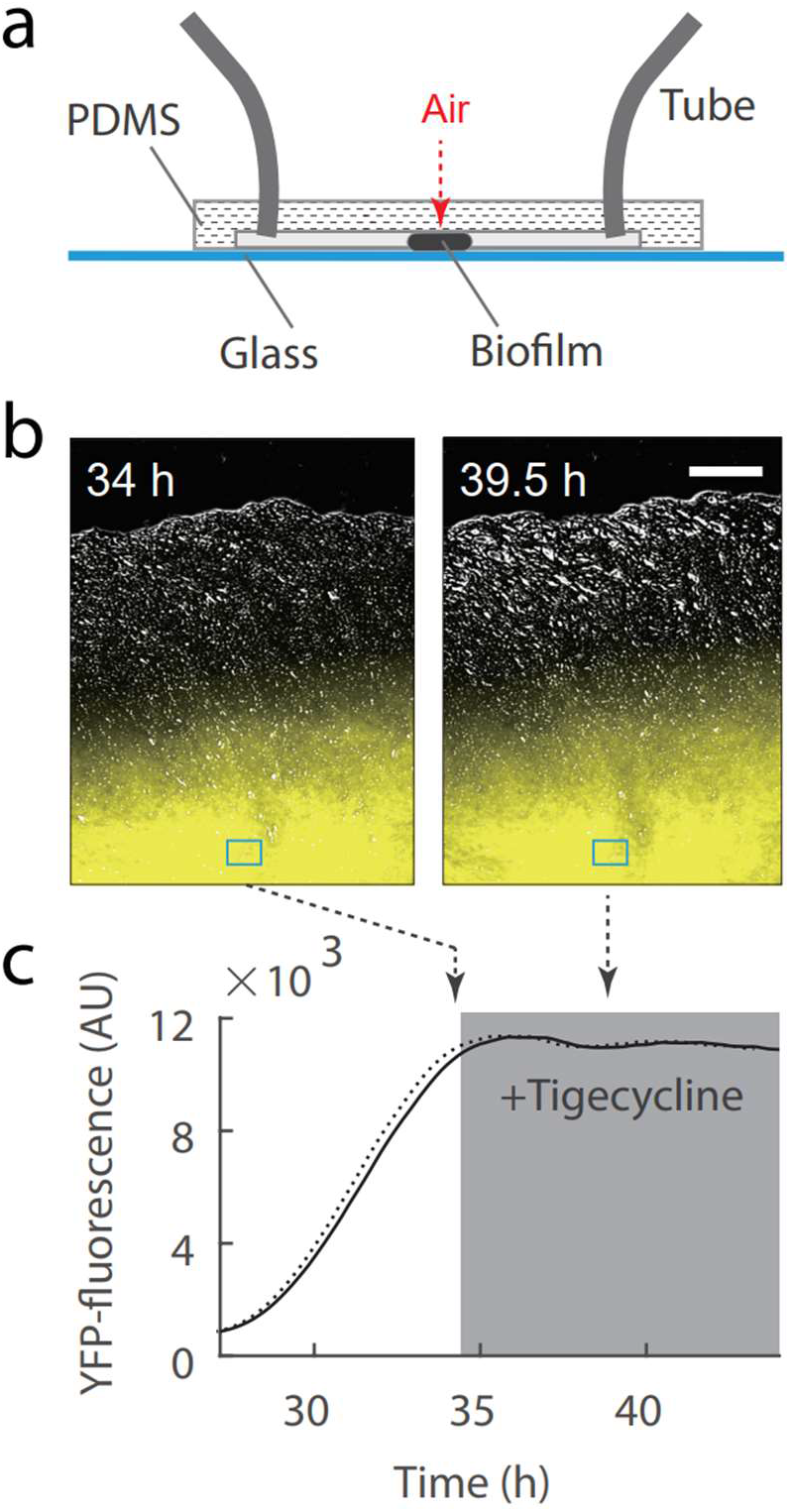
Biofilm without oxygen limitation. **a**, Side view of the microfluidic chip; The red arrow indicates how oxygen from the ambient air could diffuse through PDMS to biofilm interior. **b**, Snapshots of an *E. coli* biofilm; Composite of phase contrast and YFP (*PptsG-yfp* fluorescence, shown in yellow) channels; Scale bar, 100 μm. **c**, YFP fluorescence intensity at biofilm interior (the region marked by rectangle in **b**; The solid line shows the measured fluorescence value; The dashed line shows the estimated level of YFP protein inferred from the solid line, based on the fact that the maturation time of EYFP is 9 mins; The shaded region represents the duration of tigecycline (2 μg/ml) treatments; The dashed arrows indicate the time when the snapshots in **b** were taken.

**Supplementary Fig. 3:**
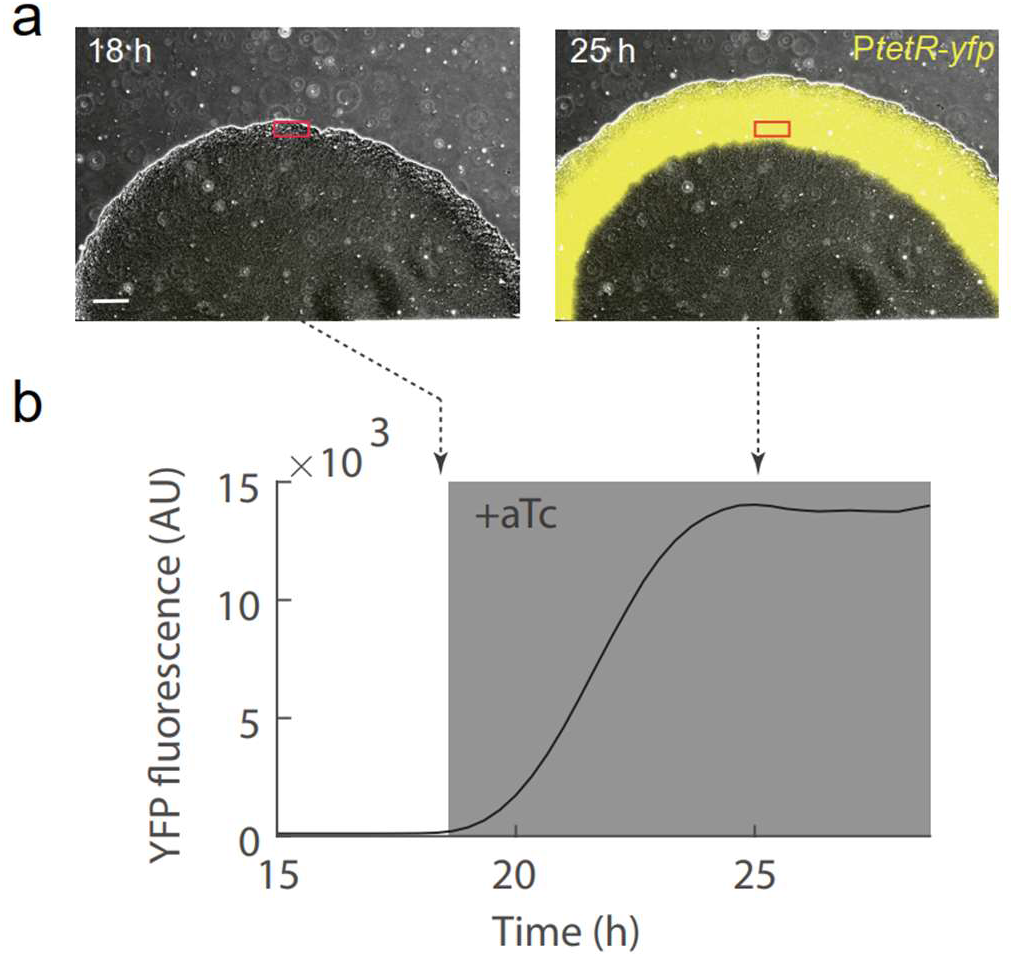
Maturation of YFP protein at biofilm periphery. **a**, Snapshots of an *E. coli* biofilm; Composite of phase contrast and YFP (*PtetR-yfp* fluorescence, shown in yellow) channels; Scale bar, 100 μm. **b**, YFP fluorescence at biofilm periphery (the region marked by rectangle in **a**); The gray shading represents the duration of aTc (anhydrotetracycline, inducer of the *tetR* promoter, 100 ng/ml) treatment; The dashed arrows indicate the time when the snapshots in **a** were taken.

**Supplementary Fig. 4:**
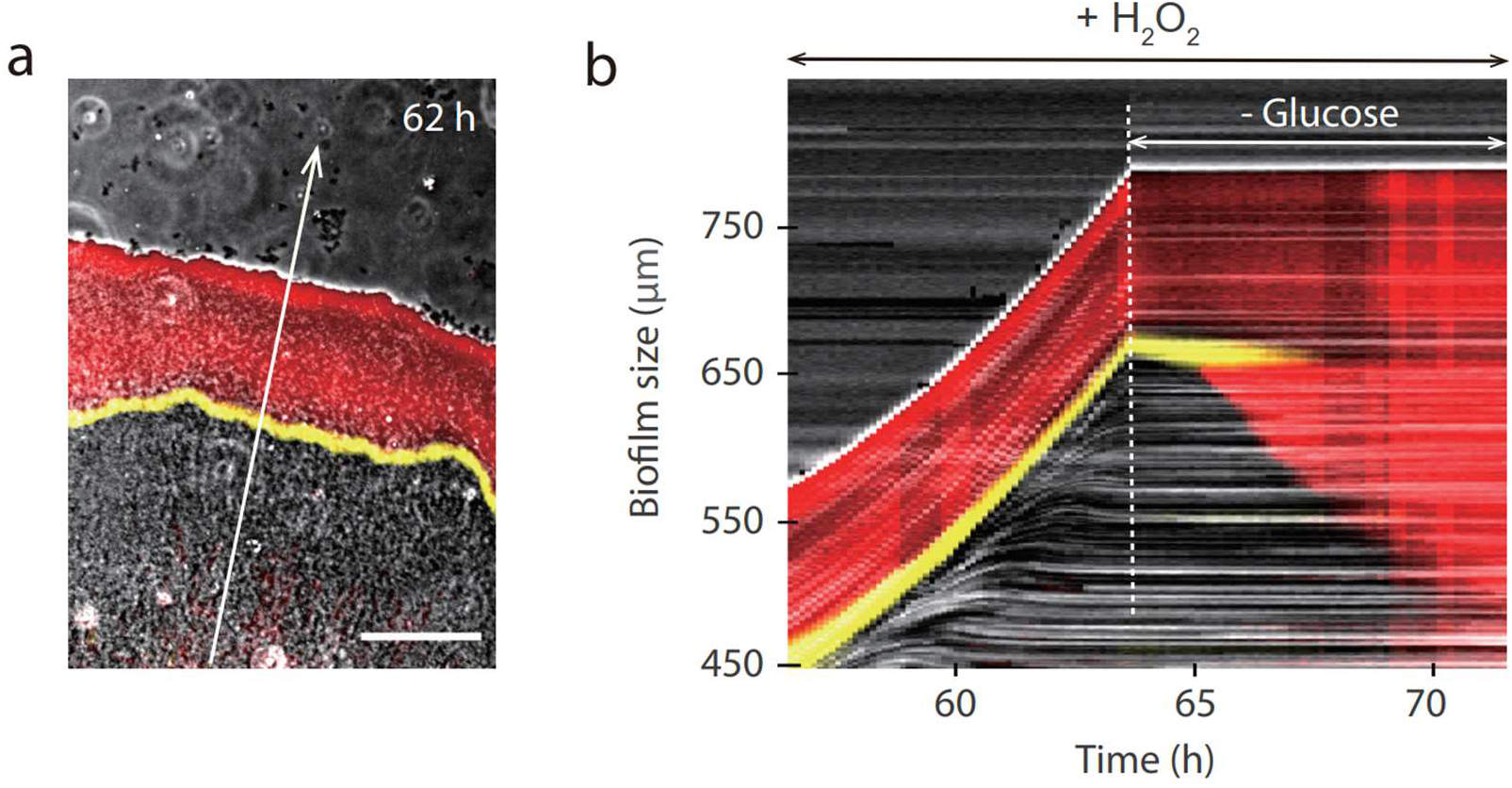
Biofilm resistance to exogeneous H_2_O_2_. Snapshot of an *E. coli* biofilm. Composite of phase contrast, YFP (DCFH-DA fluorescence, shown in red), and RFP (*PkatG-mrfp* fluorescence, shown in yellow) channels; Scale bar, 100 μm. **b**. Kymograph of biofilm response to glucose removal from the flowing medium; Measurements were taken along the line as indicated by the arrow in **a**.

**Supplementary Fig. 5:**
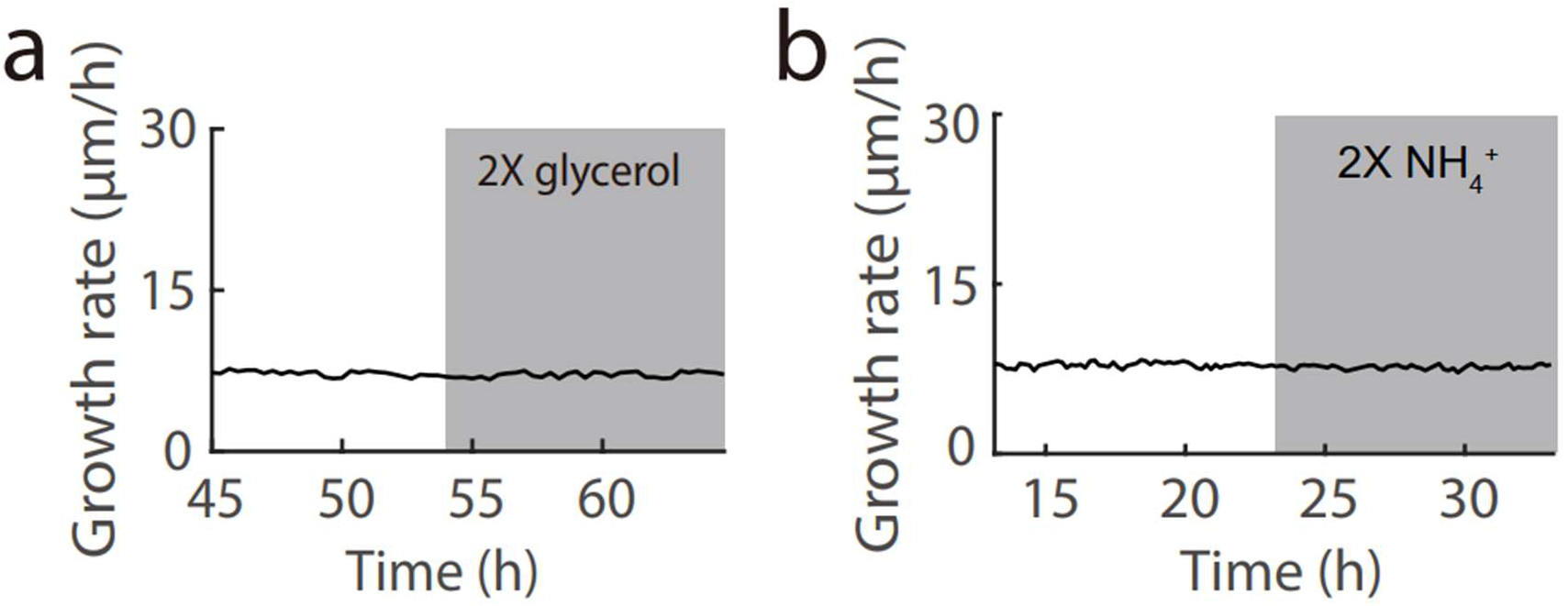
Growth of *E. coli* biofilm in modified M63B1. Glycerol (44 mM) instead of glucose was used as the carbon source. **a**, Biofilm growth rate before and after switching to 2X glycerol (88 mM). b, Biofilm growth rate before and after switching to 2X NH4^+^ (30 mM).

**Supplementary Fig. 6:**
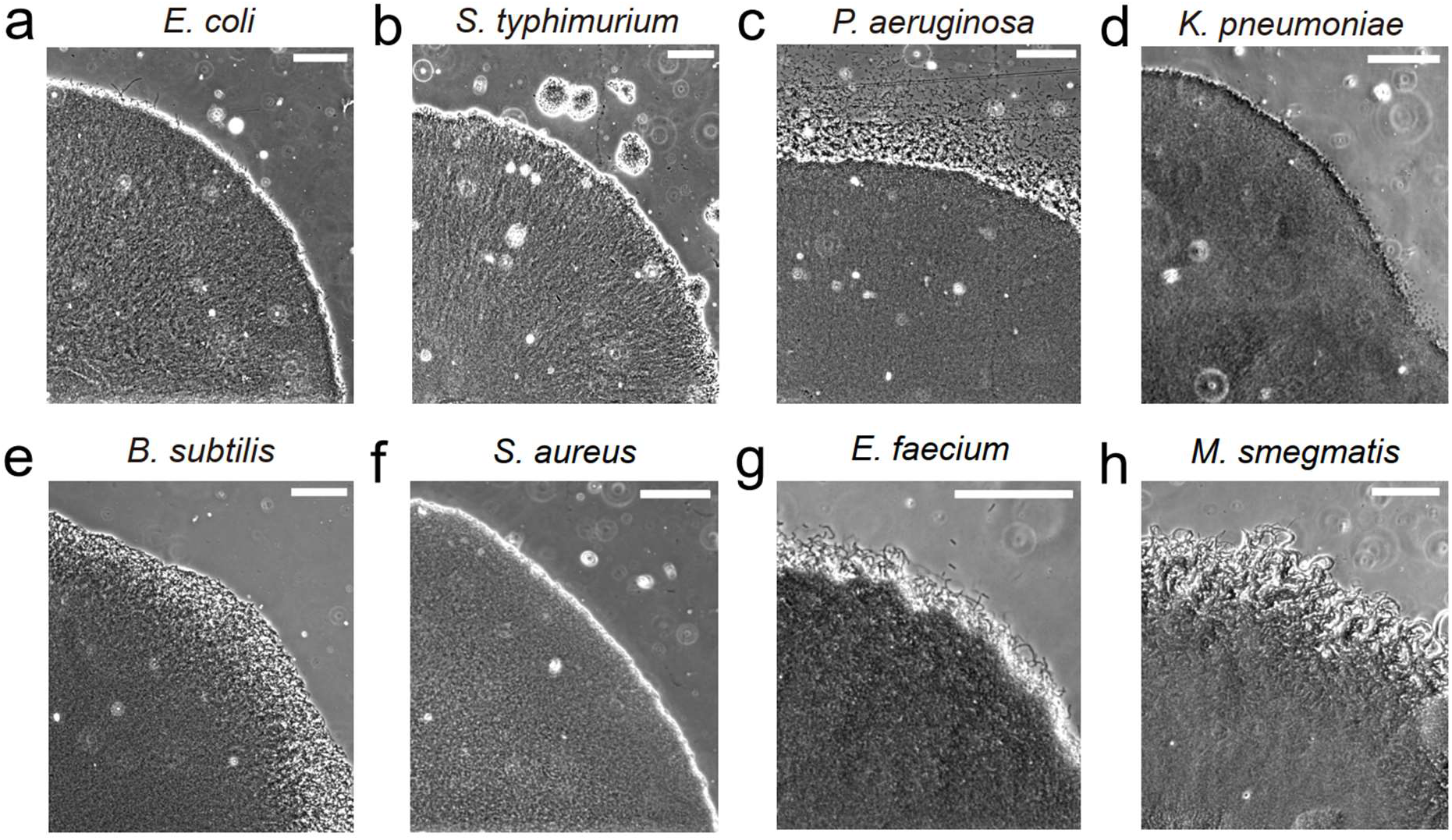
Phase contrast images of biofilms formed by commonly studied bacterial species. Scale bar, 100 μm.

**Supplementary Fig. 7:**
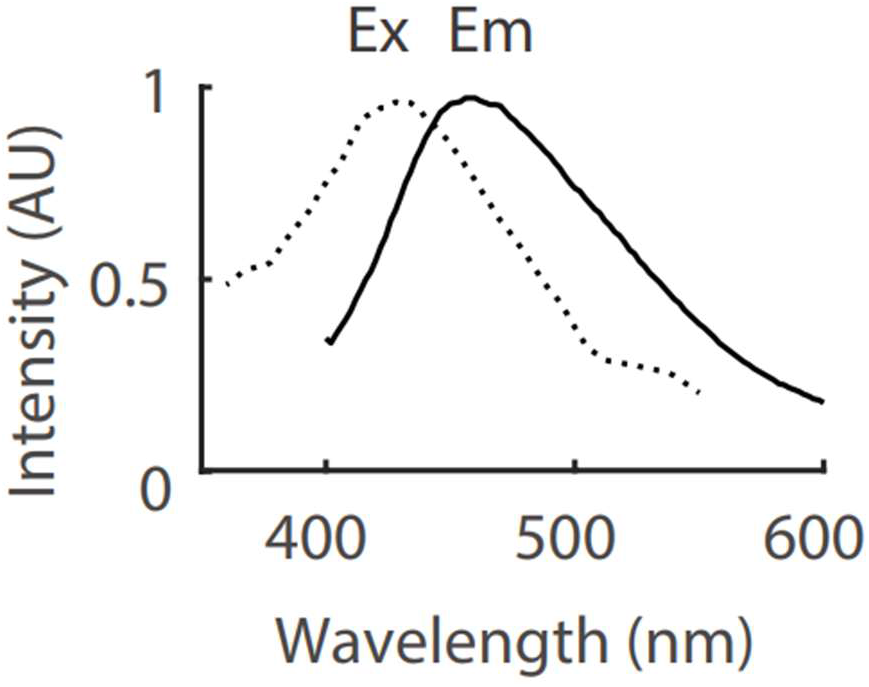
Excitation and emission spectrum of stationary phase *P. aeruginosa* culture supernatant.

**Supplementary Fig. 8:**
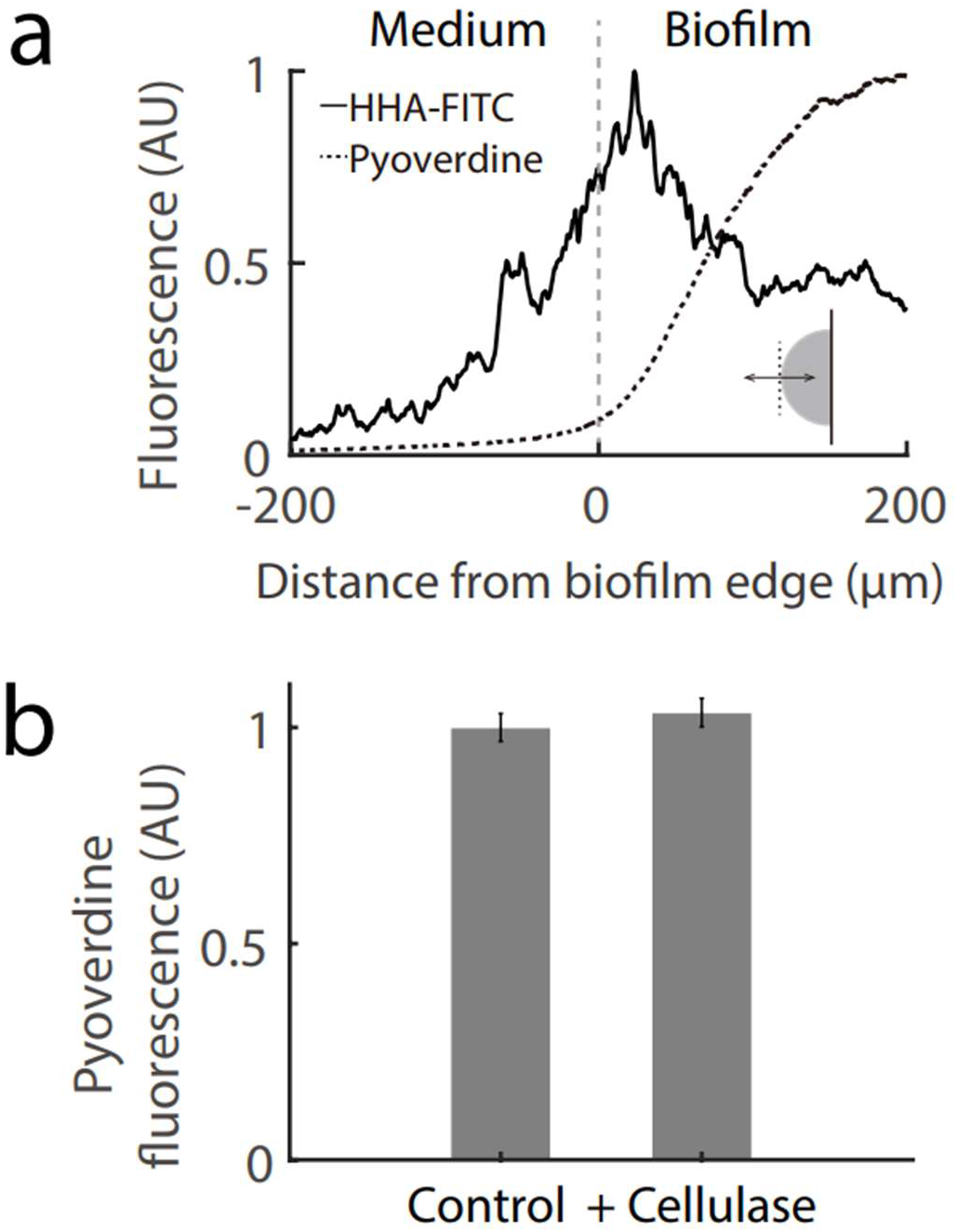
Characterization of Psl and pyoverdine. **a**, Spatial profile of HHA-FITC and pyoverdine fluorescence over a *P. aeruginosa* biofilm; The double arrow in the inset illustrates the line along which the profile was measured. **b**, Effect of cellulase addition (84 U/ml) on pyoverdine fluorescence in stationary phase *P. aeruginosa* culture. The error bars represent standard deviation, n = 3 biological replicates.

**Supplementary Fig. 9:**
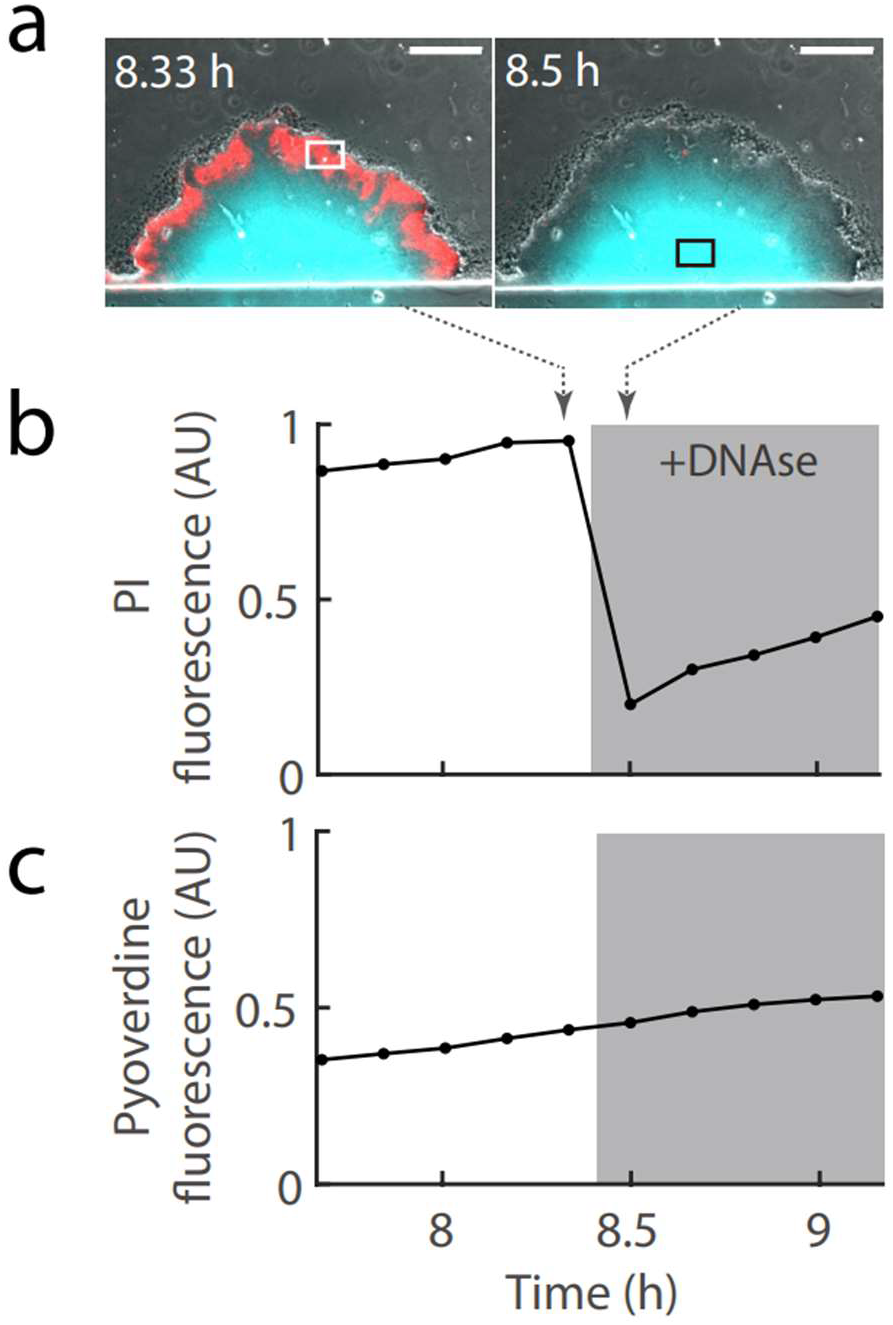
Treatment of *P. aeruginosa* biofilm with DNase. **a**, Snapshots of the biofilm; Composite of phase contrast, CFP (pyoverdine fluorescence, shown in cyan), and RFP (PI – propidium Iodide fluorescence, shown in red) channels; Scale bar, 100 μm. **b**, PI fluorescence at biofilm periphery as indicated by the white rectangle in **a**; The gray shading indicates duration of DNase treatment (750 U/ml); The dashed arrows indicate the time when the snapshots in **a** were taken. **c**, Pyoverdine fluorescence at biofilm interior as indicated by the black rectangle in **a**.

**Supplementary Fig. 10:**
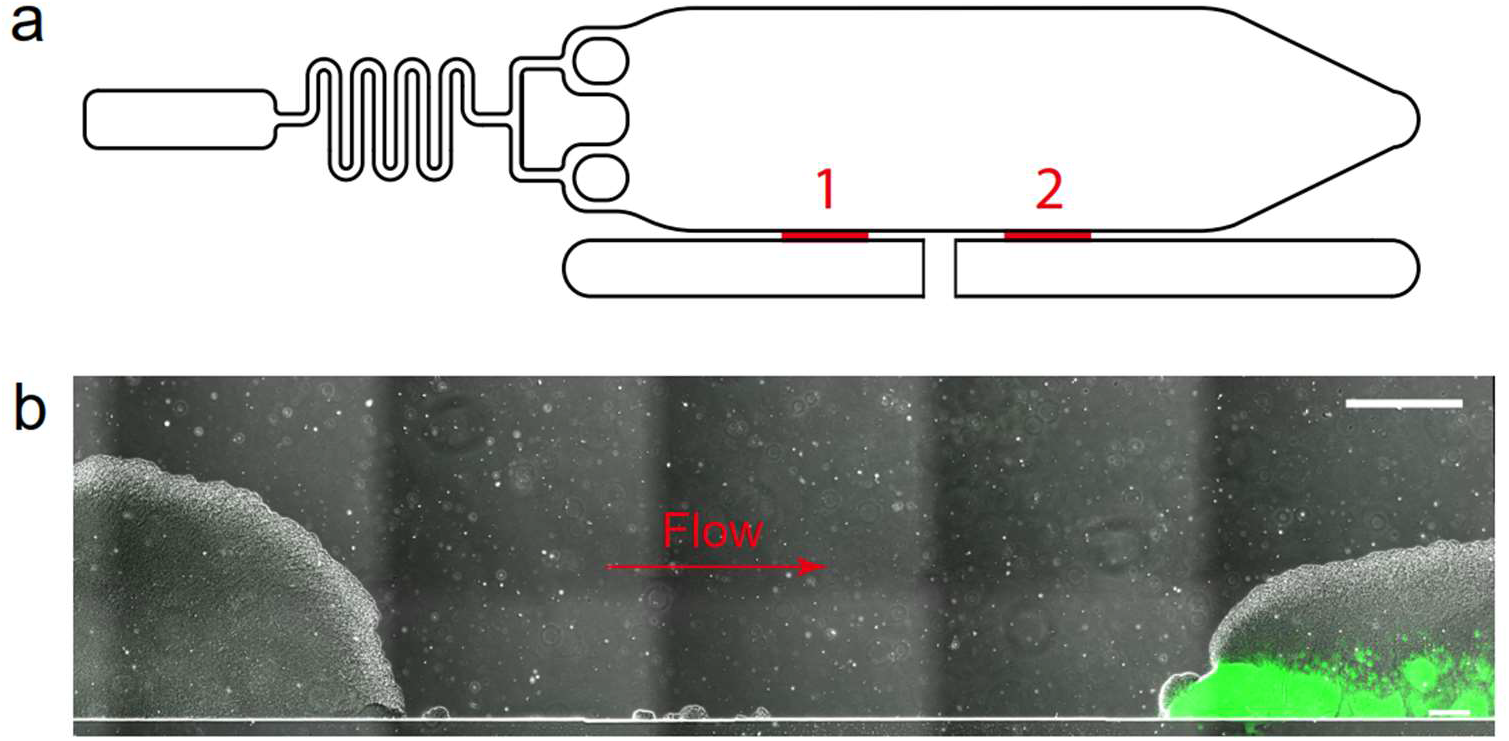
Co-culture of biofilms formed by different bacterial species. **a**, Schematic diagram of the microfluidic chip; The bacteria seeding zones are marked in red. **b**, Snapshot of two biofilms formed by *E. coli* and *P. aeruginosa*; Composite of phase contrast and GFP (indicates *P. aeruginosa*, shown in green) channels; *E. coli* and *P. aeruginosa* were loaded to the seeding zones 1 and 2 respectively, and cultured with the M63B1 medium; Scale bar, 300 μm.

**Supplementary Fig. 11:**
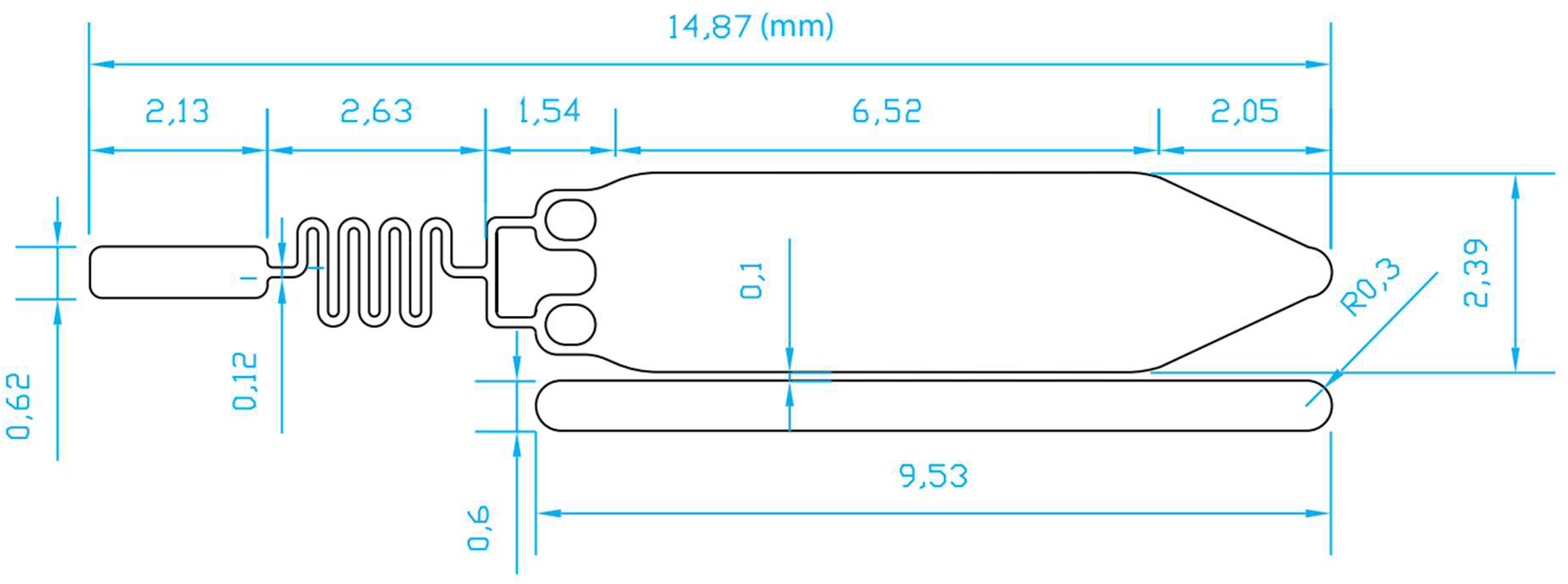
Dimensions of the microfluidic chip introduced in Fig. 1a.

**Supplementary Video 1: Time-lapse video of an *E. coli* biofilm.**

**Supplementary Video 2: Time-lapse video of a *P. aeruginosa* biofilm.** Composite of phase contrast and CFP (shown in cyan) channels;

**Supplementary Video 3: Time-lapse video of a P. *aeruginosa* biofilm.** Composite of phase contrast, CFP (pyoverdine fluorescence, shown in cyan), and GFP (HHA-FITC fluorescence, shown in red) channels; Switched to medium containing cellulose (84 U/ml) after 7h 20m.

**Supplementary Table 1:** Bacterial Strains used in this study

| Strain | Description |
| --- | --- |
| *E. coli* | BW25113 |
| *E. coli* with plasmid, pDL30 *PptsG-yfp* | *PptsG* was activated by glucose limited |
| *E. coli* with plasmid, p15A *PkatG-rfp* | *PkatG* was activated by H<sub>2</sub>O<sub>2</sub> |
| *E. coli* with plasmid, pSB1C3 *PtetR-yfp* | *PtetR* was induced by aTc |
| *Pseudomonas aeruginosa* | PAO1 |
| *Pseudomonas aeruginosa* Δ*paaP* | Less prone to dispersion in our experiments |
| *Pseudomonas aeruginosa* Δ*paaP* Δ*pvdA* | Deficient in pyoverdine synthesis |
| *Salmonella typhimurium* | ATCC 14028 |
| *Bacillus subtilis* | NCIB 3610 |
| *Klebsiella pneumoniae* | ATCC BAA-1144 |
| *Staphylococcus aureus* | RN4220 |
| *Enterococcus faecium* | ATCC 35667 |
| *Mycobacterium smegmatis* | mc<sup>2</sup>-155 |

